# *In silico* analysis of the Chikungunya virus and SARS-CoV-2 Macrodomain

**DOI:** 10.1101/2025.10.10.681726

**Authors:** Vikas Tiwari, Moira Rachman, Kenneth Huang, Ignacia Echeverria, Andrej Sali

## Abstract

The macrodomain of the Chikungunya virus NSP3 protein (ChikV Mac1) hydrolyzes mono-ADP ribose post-translational modifications on the human host proteins. Mutations in Mac1 reduce ChikV virulence. Thus, ChikV Mac1 is a viral drug target. While no potent ChikV Mac1 inhibitors are available, high-affinity inhibitors were developed for Mac1 in Severe Acute Respiratory Syndrome Coronavirus 2 (SARS-CoV-2). Here, we rationalize this difference in the ligand binding affinity in terms of three differences in the binding site structure and dynamics. First, the apo ChikV Mac1 binding site exhibits substantially more conformational heterogeneity than that of SARS-CoV-2, based on microsecond-scale molecular dynamics simulations; it also has a less druggable binding site, based on program SiteMap. Second, water binding sites overlapping with the ligand binding site in ChikV Mac1 are predicted by program WaterMap to have a stronger affinity for water molecules than the corresponding sites in SARS-CoV-2 Mac1, thus decreasing ligand binding affinity. Finally, a smaller number of Mac1 residues interacts persistently with ADP-ribose in holo ChikV Mac1 than in SARS-CoV-2. With these rationalizations in hand, we designed ligands of ChikV Mac1 using a fragment growth strategy; subsequent molecular dynamics simulations of a representative ligand in complex with ChikV Mac1 substantiated our design.

## Introduction

The Chikungunya virus (ChikV) was first isolated during an arthritic disease outbreak in Tanzania in 1952^1,2^. ChikV is a mosquito-borne virus that belongs to the alphavirus genus of the Togaviridae family. ChikV infections have emerged as a global health risk with approximately 16.9 million cases per year^3^. Major symptoms of ChikV infection include severe fever, rashes, and joint pain. Chronic arthritis-like symptoms may persist and can be debilitating^4,5^. The ChikV genome (positive-sense RNA) encodes five structural proteins and four nonstructural proteins (NSP1-4)^6^. NSP3 consists of a conserved macrodomain (Mac1) at the N-terminus, a poorly conserved hypervariable domain (HVD), and a central zinc-binding domain known as alphavirus unique domain (AUD)^7^. The macrodomain fold is highly conserved in evolution; it has been identified in bacteria, algae, and eukaryotes^8,9^. It has been suggested that ChikV Mac1 suppresses the host immune response through its ADP-ribosyl hydrolase activity^10^, which removes ADP-ribose post-translational modifications from the target host proteins by hydrolyzing mono ADP-ribosylated aspartate and glutamate residues. Thus, Mac1 has been a therapeutic target for antiviral drug discovery^10^, supported by its crucial role in ChikV virulence in mice. Despite its therapeutic potential, efforts to identify ChikV inhibitors have had limited success. A fragment screen of ∼14,000 compounds identified only weak inhibitors (*e*.*g*., 2-pyrimidone-4-carboxylic acid scaffold)^11^. Another computational docking and simulation study screened 820 compounds and predicted that natural compounds from plants, including Apigetrin, Baicalin, Baloxavir, Luteoloside, Rutaecarpine, and Amentoflavone^12^, were Mac1 inhibitors. Another study identified N-[2-(5-methoxy-1H-indol-3-yl) ethyl]-2-oxo-1,2-dihydroquinoline-4-carboxamide through virtual screening of 245,532 natural compounds, followed by *in vitro* validation through a microscale thermophoresis binding assay and *in vivo* inhibition of ChikV replication^13^.

Similar to ChikV, SARS-CoV-2 NSP3 contains three tandem macrodomains, with Mac1 serving as the catalytically active macrodomain that binds and hydrolyzes mono-ADP-ribose on posttranslationally modified target host proteins^14,15^. SARS-CoV-2 Mac1 is essential for viral pathogenesis and represents a promising drug target^16,17^. In contrast to ChikV Mac1, it has proven amenable to inhibitor development. An early crystallographic screen of approximately 2,600 compounds revealed 234 fragment structures bound to SARS-CoV-2 Mac1^18^. Using these hits, several optimized inhibitors were designed, followed by another round of crystallographic screening^19^. Among the resulting top inhibitors was AVI-4206, a potent inhibitor with an IC_50_ of 20nM that is effective in an animal model of SARS-CoV-2 infection^20^. Other studies have identified additional promising scaffolds, including 2-amide-3-methylester thiophene scaffold derivatives that bind SARS-CoV-2 Mac1 (IC_50_ = 1.5 μM) and inhibit viral replication^21^, synthetic analogs of ADP-ribose that bind SARS-CoV-2 Mac1 with nanomolar affinity^22^, and pyrrolo-pyrimidine-based compounds that inhibit viral replication in SARS-CoV-2^23^.

In contrast to SARS-CoV-2, ChikV Mac1 has remained a difficult target to inhibit, despite its structural similarity to SARS-CoV-2 Mac1^24^. Here, we hypothesized that characterizing structural dynamics of ChikV Mac1 can help (i) explain why this protein is more difficult to target than SARS-CoV-2 Mac1 and (ii) design more potent inhibitors. Thus, we proceeded by performing long molecular dynamics simulations of various forms of Mac1, followed by analyzing the resulting conformations in terms of druggability and interactions with water molecules and potential inhibitors. Armed with the resulting rationalizations, we proposed several inhibitor designs and characterized them with additional molecular dynamics simulations.

## Methods

### Sequence comparison and evolutionary analysis

ChikV and SARS-CoV-2 Mac1 sequences from their structure files deposited in the Protein Data Bank (PDB)^25^ (PDB IDs 3GPO and 6W02, respectively) were aligned using EMBOSS Needle. Alignments were visualised using ESPript3^26–29^. Residue-level conservation was estimated using ConSurf, with default parameters for homolog searching (*i*.*e*., one HMMER iteration, with a maximum of 150 homologs and E-value cutoff of 0.0001), multiple sequence alignment (MAFFT-L-INS-i), and phylogenetic tree construction^30^.

### Molecular dynamics simulations

Simulations were performed using Desmond from the Schrödinger package^31^ and the OPLS4 forcefield^32^. The crystal structures of holo Mac1 bound to ADP ribose (PDB IDs 3GPO and 6W02 for ChikV and SARS-CoV-2 Mac1, respectively) and apo Mac1 (PDB IDs 3GPG and 6WEY for ChikV and SARS-CoV-2 Mac1, respectively) were used as starting points for the simulations^26,27,33^. These starting structures were first processed using the Schrödinger *protein preparation wizard* module, including protonation at a specified *p*H, hydrogen bond optimization, and restrained minimization^34^. The simulation system was set up using the Schrödinger *system builder* module. Prepared structures were solvated using the TIP3P water model in an orthorhombic box. The system was then neutralized by adding counter ions; an ionic concentration of 0.15 M was established using NaCl. The final system was equilibrated using the default relaxation protocol that includes a series of short simulations: (1) Brownian dynamics simulations of the NVT ensemble at 10 K for 100 ps with restraints on solute non-hydrogen atoms, (2) NVT molecular dynamics simulations for 12 ps at 10 K with restraints on solute non-hydrogen atoms, (3) NPT molecular dynamics simulations at 10K for 12 ps with restraints on solute non-hydrogen atoms, (4) NPT molecular dynamics simulations for 12 ps with restraints on solute non-hydrogen atoms, and (5) NPT molecular dynamics simulations for 24 ps without restraints. Production simulations were performed in the NPT ensemble for 1000 ns. Three independent runs were executed with different initial seeds for ChikV (apo), SARS-CoV-2 (apo), and their ADP-ribose-bound counterparts.

The molecular dynamics trajectories were analyzed using the *Simulation Interaction Diagram (SID)* module of Desmond. Root-mean-square deviation (RMSD) was calculated for each frame using the initial frame as the reference. The ligand RMSD indicates the RMSD of ligand non-hydrogen atoms calculated after superposing the protein-ligand complex onto the protein backbone of the reference frame. Higher values of the ligand *vs* protein RMSD imply ligand diffusion away from its initial binding site. Root-Mean-Square Fluctuation (RMSF) was calculated for each residue to quantify local changes. Non-covalent interactions (*i*.*e*., hydrogen bonds, hydrophobic interactions, ionic interactions, and water bridges) between protein and ligand were monitored throughout the simulations. The consensus residues were defined as residues that interact with ADP-ribose in all three trajectory replicates in more than 30% of the snapshots. Binding energies of ADP-ribose were calculated across the trajectories using the Schrödinger “thermal_mmgbsa.py” script. Binding site analysis of apo structures across trajectories was done using the Schrödinger “trajectory_binding_site_volumes.py” script. The SiteMap parameters were optimized to identify the binding site across all sampled conformational states: sitebox size was set to 10 Å, the enclosure value to 0.3, and the merging distance threshold was set to 10 Å. The SiteMap analysis provides three key metrics for evaluating binding sites: volume, site-score, and druggability score (Dscore); higher Dscore values indicate better druggability potential.

### WaterMap analysis

WaterMap calculations were performed on both apo and holo Mac1 structures^35,36^. The default parameters were used, with water molecules within 10 Å of the ligand considered for the holo Mac1 simulations. For apo structures, the ADP-ribose binding residues (residues within 4 Å of ADP-ribose) were specified to define the analysis region. Simulations were performed for 2 ns using the OPLS4 force field and TIP4P water model. In each case, three replicates were performed using different random seed values.

### Ligand design by fragment growth

We designed inhibitors using a fragment-based approach, relying on the recent deposition of structures of ∼100 fragments (typically, less than 100 Da in mass) bound to ChikV Mac1, determined by X-ray crystallography^24^. 1,730 small molecule analogs of these fragments (typically, 100-200 Da in mass) were downloaded from http://arthor.docking.org, containing the structure of a starting fragment from the PDB structure (PDB ID 7H8P). This fragment was chosen as an initial starting structure for the inhibitor design due to the large number of available analogs and correlation with stable interactions (Asp10, Ile11) identified in the molecular dynamics simulations (**Figure 3**). The analog structures were prepared with Ligprep and docked with Glide (grids were prepared with Maestro) while constraining the fragment (<0.5Å)^37–40^. 596 of the 1,730 analogs passed a score threshold of −5 (only the best-scoring isomer was considered). These 596 compounds were further filtered by requiring (i) a hydrogen bond to Asp10, Ile11, and at least one other residue; (ii) 2 or less rotatable bonds (to minimize entropic penalty for binding), (iii) 2 or less chiral centers (to minimize artefacts from experimental assays); and (iv) no non-interacting polar hydrogens (to minimize desolvation penalty for binding). A final list of 65 analogs remained, of which those interacting with Val33 were considered for further analyses.

## Results and Discussion

There has been limited success in finding potent inhibitors of ChikV Mac1. In contrast, SARS-CoV-2 Mac1 has been successfully targeted through systematic screening efforts and structure-based design. We hypothesized that a comparison between ChkiV and SARS-CoV-2 Mac1 could inform drug design against ChikV Mac1. Therefore, we analyzed the structures, dynamics, and hydration sites of Mac1 to rationalize ChikV - SARS-CoV-2 Mac1 differences in ligand binding and druggability. We also applied the lessons learnt to design predicted potent ChikV Mac inhibitors.

### Structure analysis: The folds and ligand binding sites of Mac1 are similar in ChikV and SARS-CoV-2

We begin by quantifying the sequence conservation of ChikV Mac1, using its multiple sequence alignment with easily identifiable homologs (Methods); this alignment does not include distant homologs, such as SARS-CoV-2 Mac1. The sequence conservation score was mapped on the holo ChikV Mac1 structure. Several of the residues within 4 Å of the ADP-ribose substrate were not conserved, including Val113, Ser115, Arg144, and Trp148 (**Figure 1c**), even though Val113 backbone and Arg144 sidechain form H-bonds with ADP-ribose.

**Figure 1:**
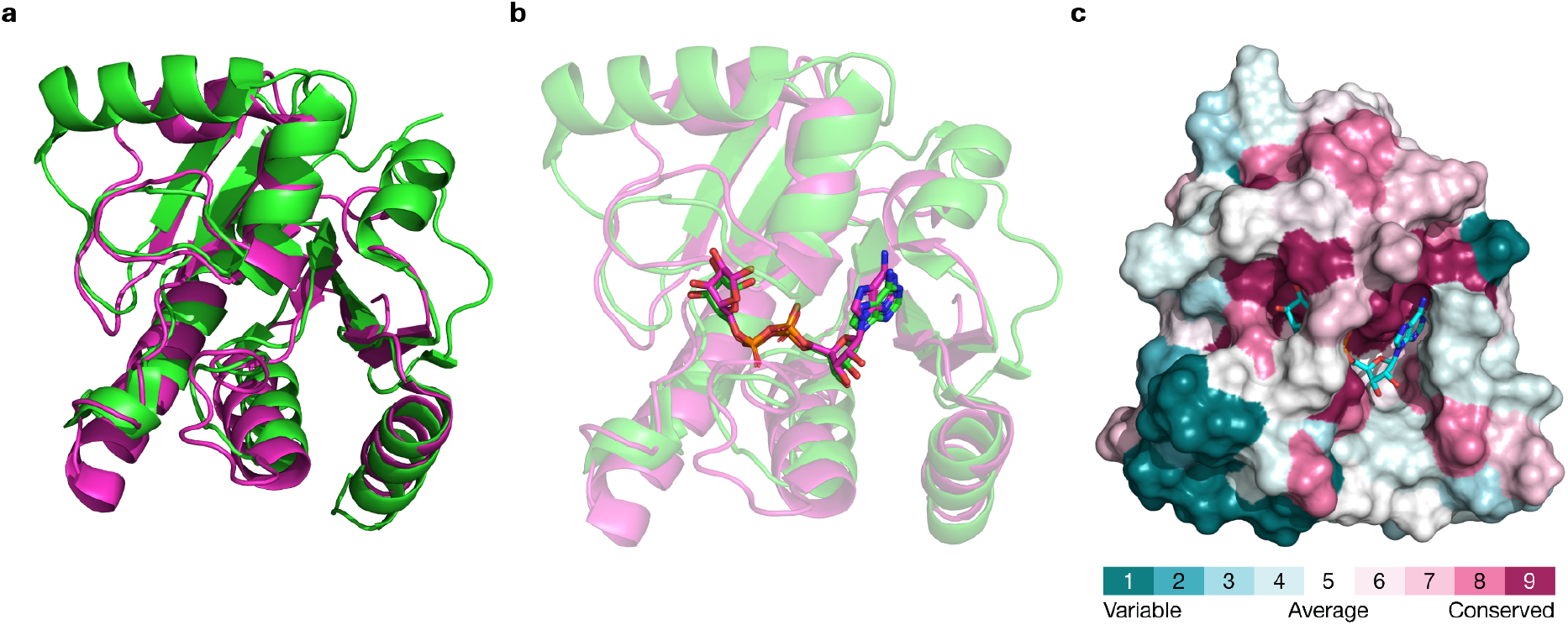
Structures and sequence divergence of ChikV Mac1. (a) Superposition of apo Mac1 of SARS-CoV-2 (green) and ChikV (magenta) (RMSD: 3.1 Å). (b) Superposition of holo Mac1 of SARS-CoV-2 (green) and ChikV (magenta) (RMSD: 2.7 Å). (c) Residue conservation of ChikV Mac1 across homologs was determined with ConSurf. The consurf scores are mapped on the apo ChikV Mac1 structure. The apo ChikV Mac1 was superposed on holo ChikV Mac1 structure and ADP-ribose (stick representation) is shown.

Next, we compared the sequences and structures of ChikV and SARS-CoV-2 Mac1. ChikV and SARS-CoV-2 Mac1 share low sequence similarity (22.7% sequence identity; **Supplementary Figure 1**). Nevertheless, the two proteins have similar structures, in both apo and holo forms (**Figure 1a**): the all-atom RMSD is 3.1 Å and 2.7 Å for the apo and holo structures, respectively (**Figure 1b**). The binding pose of the ADP-ribose substrate is also similar in both proteins, although there are several residue differences, including Thr111, Val113, Tyr114, and Arg144 in ChikV.

Because the ligand binding sites in ChikV and SARS-CoV-2 Mac1 are relatively similar, the corresponding small structural differences in the binding site do not offer an obvious rationalization of their significantly different druggabilities. Therefore, in an attempt to find such a rationalization, we evaluated their dynamics and hydration, as follows.

### Dynamics analysis: Apo Mac1 is more flexible in ChikV than SARS-CoV-2

All-atom molecular dynamics simulations in explicit solvent were performed to evaluate the dynamics of apo Mac1 in ChikV and SARS-CoV-2. Three independent simulations were conducted for each system, using different random seeds. Overall backbone RMSD values show higher conformational flexibility of apo ChikV Mac1 compared to SARS-CoV-2 (**Figure 2a**). Despite being more dynamic, ChikV Mac1 remains structurally stable throughout the simulations, as evidenced by its radius of gyration (**Supplementary Figure 2**). In agreement with these overall RMSD values, the individual residues of apo ChikV Mac1 are generally more flexible than the corresponding residues in apo SARS-CoV-2 Mac1, based on the residue RMSF values (**Figure 2b-d**); notably, ADP-ribose binding site residues also show higher flexibility in apo ChikV Mac1 than in SARS-CoV-2. In addition, apo ChikV Mac1 formed fewer persistent (>80% of simulation time) intra-protein H-bonds compared to SARS-CoV-2 (**Supplementary Figure 3**). In conclusion, apo SARS-CoV-2 Mac1 adopts a more rigid conformation than apo ChikV Mac1. This finding raises the possibility that ligand binding to ChikV Mac1 is accompanied by a larger entropic penalty than ligand binding to a more rigid SARS-CoV-2 Mac1.

**Figure 2:**
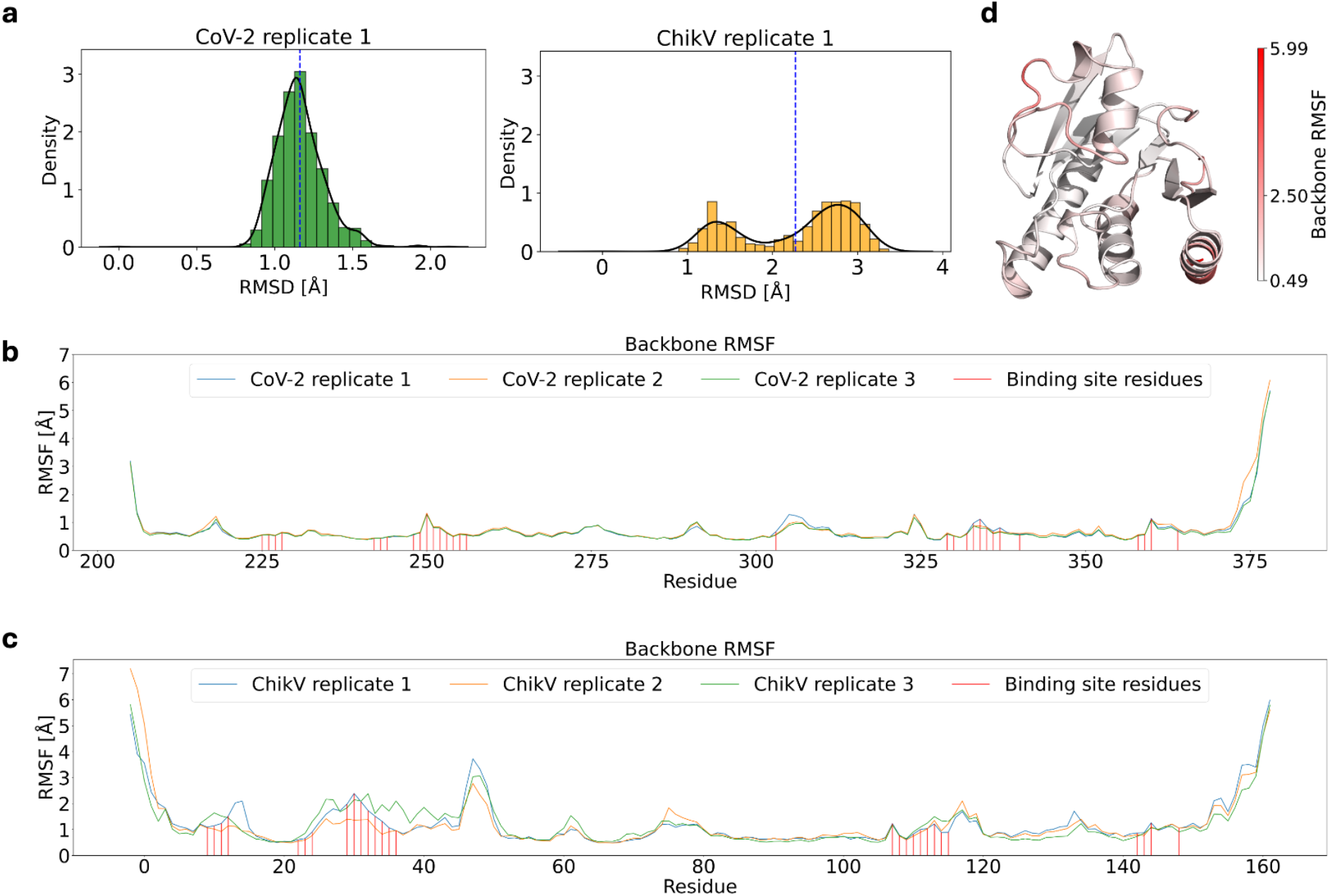
Analysis of structural dynamics of apo SARS-CoV-2 and ChikV Mac1 from MD simulations. (a) RMSD plot with the initial structure as the reference structure. (b, c) Root-mean-square fluctuation (RMSF) per residue for the backbone. (d) RMSF mapped on the structure of ChikV Mac1.

Further, to assess the druggability of the ADP-ribose binding site, we used the SiteMap tool of Schrödinger to analyse the binding site characteristics across all simulation frames. Initial analysis using default SiteMap parameters failed to consistently predict the binding site in some of the conformational states. Thus, the parameters were optimised to ensure reliable binding site predictions across all the simulation frames (Methods). The ChikV Mac1 shows a lower mean site-score and D-score than SARS-CoV-2 Mac1 (**Supplementary Figure 4**). These scores indicate a lower druggability of the ChikV Mac1 binding site compared to that of SARS-CoV-2 Mac1, consistent with the relative dearth of ChikV Mac1 inhibitors.

### Dynamics analysis: ADP-ribose interaction is more dynamic in ChikV than SARS-CoV-2 Mac1

We performed molecular dynamics simulations of these two complexes, followed by three comparisons of simulated ADP-ribose - Mac1 interactions between the two species and apo/holo states.

First, the ribose ring was observed to be more dynamic in ChikV than SARS-CoV-2 Mac1 (**Figure 3a, Supplementary Movie**). Specifically, in ChikV Mac1, consensus interacting residues were Asp10, Ile11, and Val33, based on three simulation replicates (Methods; **Supplementary Figures 5-7**). In contrast, SARS-CoV-2 Mac1 had a larger number of consensus residues, including Asp22, Ile23, Val49, Ala50, Ser128, Ala129, Ile131, Phe132, and Phe156 (**Supplementary Figures 8-10**). In addition, the magnitude of estimated ADP-ribose - ChikV Mac1 binding energy (Methods) decreased during the course of simulation. In contrast, in SARS-CoV-2 Mac1, it remained stable in all replicates (**Figure 3d and Supplementary Figure 11**). Second, the ligand binding site becomes more rigid after ADP-ribose binding in both species (**Supplementary Figure 12**). Specifically, ADP-ribose binding reduced the mean backbone RMSF of all binding site residues in both species, except for Cys143 and Arg144 in ChikV Mac1. Third, ADP-ribose rigidifies its binding site more in SARS-CoV-2 than ChikV (**Supplementary Figure 12**). In addition, the ADP-ribose binding site is more rigid in SARS-CoV-2 than ChikV, for both apo and holo states.

**Figure 3:**
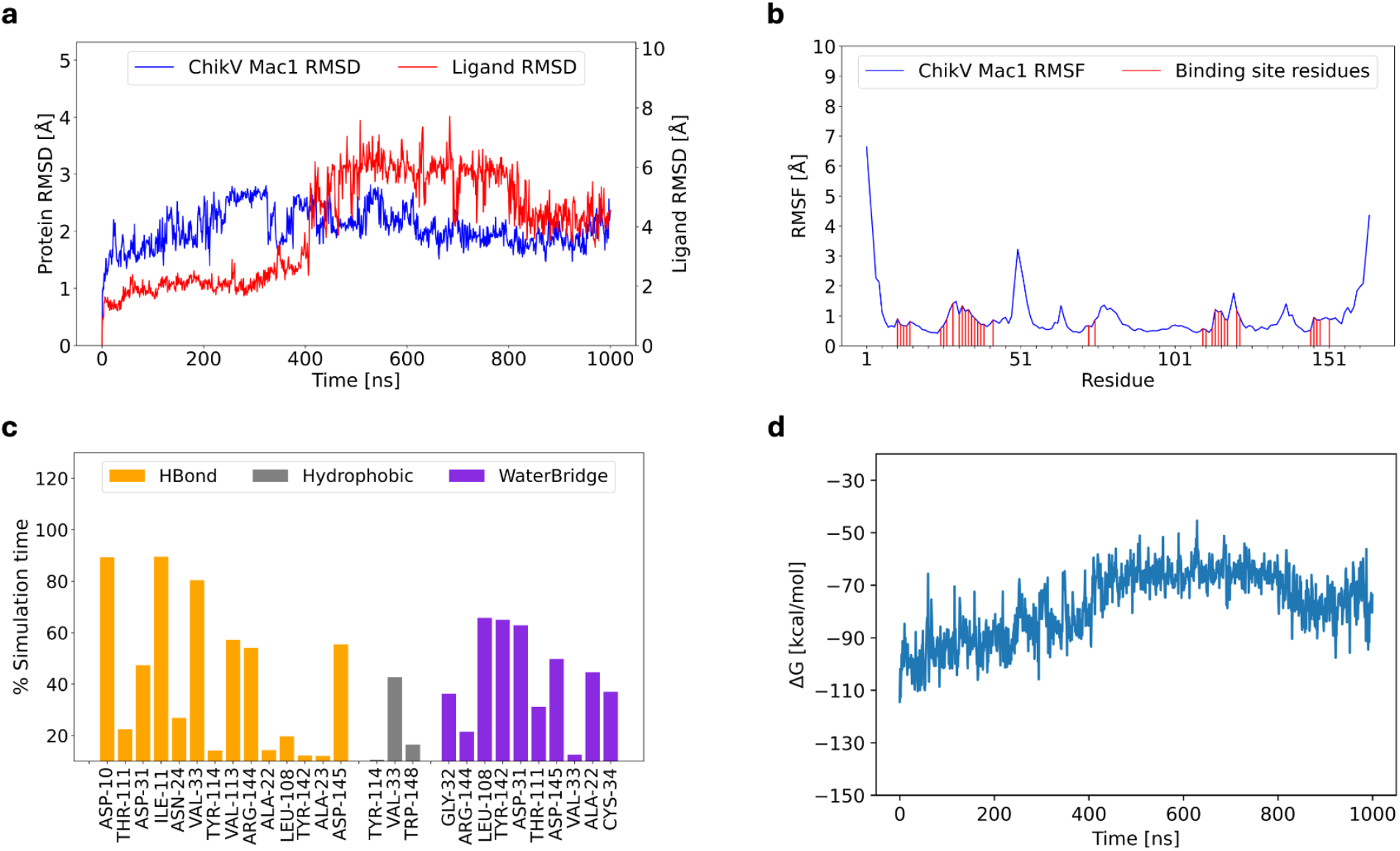
Molecular dynamics simulation of holo ChikV Mac1. (a) RMSD plot of Mac1 and ADP-ribose (ligand RMSD). (b) RMSF of ChikV Mac1. The ADP-ribose interacting residues are marked. (c) Non-covalent interaction between ADP-ribose and ChikV Mac1. The y-axis represents the persistence of each interaction type. Transient interactions (<10% persistence) have not been shown. (d) The binding energy (ΔG) of ADP-ribose across the simulation time.

In conclusion, these three simulation comparisons help rationalize the experimentally observed preference of ADP-ribose for SARS-CoV-2 Mac1 rather than ChikV Mac1. Moreover, this analysis suggests specific ligand interactions for consideration in both the design of high-affinity ChikV Mac1 ligands and filtering ligands identified by virtual screening.

### Hydration site analysis: Hydration is less unfavorable for ligand binding in SARS-CoV-2 than ChikV Mac1

Interactions of water molecules with ligand and/or protein atoms can influence the ligand binding affinity. In particular, a water molecule can compete with ligand binding or it can serve as a bridge between the ligand and protein. In the former case, the weaker the water molecule binds to the protein, the less unfavorable (*i*.*e*., relatively more favorable) it is for ligand binding.

To study the hydration of Mac1, we used the WaterMap program^35^. WaterMap employs short molecular dynamics simulations to cluster water molecules into hydration sites around a defined surface patch, such as the ADP-ribose binding site, followed by calculating enthalpy, entropy, and free energy of water binding for each hydration site. A hydration site is defined by the coordinates of the water molecules in the corresponding cluster. A water molecule in an “unstable” hydration site (ΔG > 0 kcal/mol) can be either “replaced” or “displaced” by a ligand^41,42^; a water molecule is said to be replaced when ΔH < 0 kcal/mol, often corresponding to ligand-protein interactions that are similar to water-protein interactions, while a water molecule is said to be displaced when ΔH > 0 kcal/mol. Therefore, displacement of unstable hydration sites is more likely to be favorable for ligand binding than displacement of stable hydration sites^43^. We focus here on the hydration sites with at least one water molecule within 4 Å of any atom of any of the ligand binding residues in the holo structure (which in turn are defined as residues with at least one atom within 4 Å of any ligand atom); ligand binding residues in an apo structure are defined based on the alignment with the corresponding holo structure. Apo Mac1 in ChikV and SARS-CoV-2 both have 111 such hydration sites.

The apo ChikV Mac1 has fewer displaceable hydration sites and a larger number of stable hydration sites than apo SARS-CoV-2 Mac1, across all WaterMap replicates (**Figure 4a, 4b; Table 1**; **Supplementary Figure 14**). Specifically, the Watermap replicates average in 3.3 less displaceable and 2.6 more stable hydration sites in apo Mac1 in ChikV than SARS-CoV-2 (**Supplementary Figure 14**). These observations suggest that the binding pocket of apo ChikV Mac1 is less druggable than in SARS-CoV-2. A similar WaterMap analysis was also performed for holo Mac1 of both species. Similarly to apo Mac1, ADP-ribose binding displaced more unstable hydration sites in SARS-CoV-2 than ChikV Mac1 (**Figure 4c, 4d**).

**Table 1:** WaterMap analysis of apo Mac1 in ChikV and SARS-COV-2.

| Hydration site properties | Apo ChikV Mac1 | Apo SARS-COV-2 Mac1 |
| --- | --- | --- |
| Number of sites within 4Å | 111 | 111 |
| $\Delta G$ range [kcal/mol] | -1.9 to 9.2 | -1.3 to 8.4 |
| Number of unstable hydration sites | 94 | 96 |
| Number of displaceable hydration sites | 38 | 46 |
| Number of replaceable hydration sites | 56 | 48 |
| Number of stable hydration sites | 17 | 15 |

**Figure 4:**
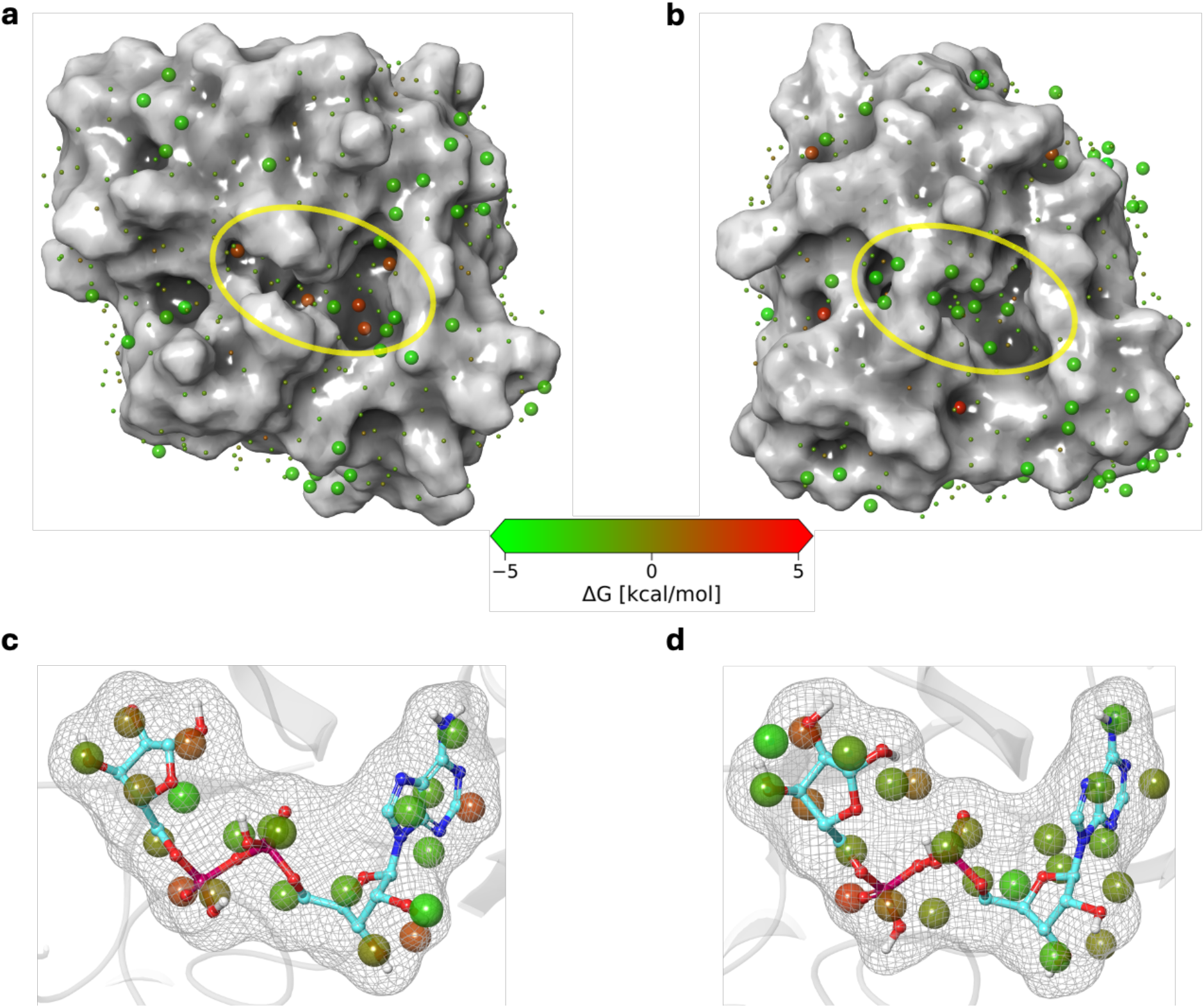
WaterMap analysis of apo and holo Mac1. (a) SARS-CoV-2 Mac1 hydration sites. (b) ChikV Mac1 hydration sites. (c) Holo SARS-CoV-2 Mac1 showing hydration sites that will be replaced upon binding of ADP-ribose. (d) Holo ChikV Mac1 showing hydration sites that will be replaced upon binding of ADP-ribose.

To evaluate if the hydration site affinity can be improved through targeted mutation, we analyzed in detail a sample hydration site in holo ChikV Mac1, near the purine ring of ADP-ribose. This hydration site is predicted to be replaceable (ΔG = 2.1 kcal/mol; ΔH = −0.7 kcal/mol). The water molecules in this site interact with Arg144 through an H-bond. In contrast to ChikV, the equivalent SARS-CoV-2 hydration site is displaceable (ΔG = 5.1 kcal/mol; ΔH = 1.4 kcal/mol). *In silico* mutation of ChikV Mac1 Arg144 to Phe (as found in SARS-CoV-2) also results in a displaceable hydration site (ΔG = 3.4 kcal/mol; ΔH = 0.1 kcal/mol). Thus, we hypothesize that the Arg144Phe point mutant of ChikV Mac1 is more druggable than the wild type, highlighting the relatively poor druggability of wild-type ChikV Mac1.

In conclusion, the hydration site analysis helps explain why the ChikV Mac1 binding pocket is a more difficult drug target than the SARS-CoV-2 Mac1.

### Guidelines for ligand design

The above analyses inspired the following two guidelines for designing potent inhibitors of ChikV Mac1:

First, ligands that can establish interactions with the consensus residues (Asp10, Ile11, and Val33) should be prioritized. This guideline is based on the evaluation of ADP-ribose interactions with ChikV Mac1 across molecular dynamics trajectories (**Figure 3**). These consensus residues show high sequence conservation among the ChikV Mac1 homologs (**Figure 1c**) as well as with SARS-CoV-2 Mac1.

Second, ligands that can displace as many of the 38 displaceable hydration sites predicted for apo ChikV Mac1 as possible should be prioritized. This guideline is based on the evaluation of hydration sites of apo and holo ChikV Mac1 (**Figure 4)**. WaterMap analysis may also help in predicting the direction in which the ligand can be grown to improve its binding affinity.

### Application of ligand design guidelines to ChikV Mac1

With the two guidelines in hand, we proceeded with predicting new ligands of ChikV Mac1, using the fragment growth approach (Methods). A previously identified fragment that interacted with Asp10 and Ile11^24^ was grown to interact with Val33, per the first guideline. One of the top hit compounds (x0190) was selected for further analysis. First, the interaction stability of x0190 with ChikV Mac1 was assessed through molecular dynamics simulations. x0190 remained stable throughout the simulated trajectory, interacting with Ile11, Val33, Arg144, and Trp148 of ChikV Mac1, at least partly satisfying the first guideline. Similar behavior was observed in two additional replica trajectories (**Supplementary Figures 15-17**).

Next, to further optimize x0190, we considered the second guideline, starting with a WaterMap analysis of the predicted x0190 binding site. WaterMap predicted unstable hydration sites near the ribose-binding site in Mac1 (**Figure 5**). Thus, we hypothesized that x0190 can be further grown using the analogue-by-catalogue approach. For example, addition of a pyridine ring may displace additional unstable hydration sites while simultaneously establishing productive interactions with Tyr114, thereby improving binding affinity.

**Figure 5:**
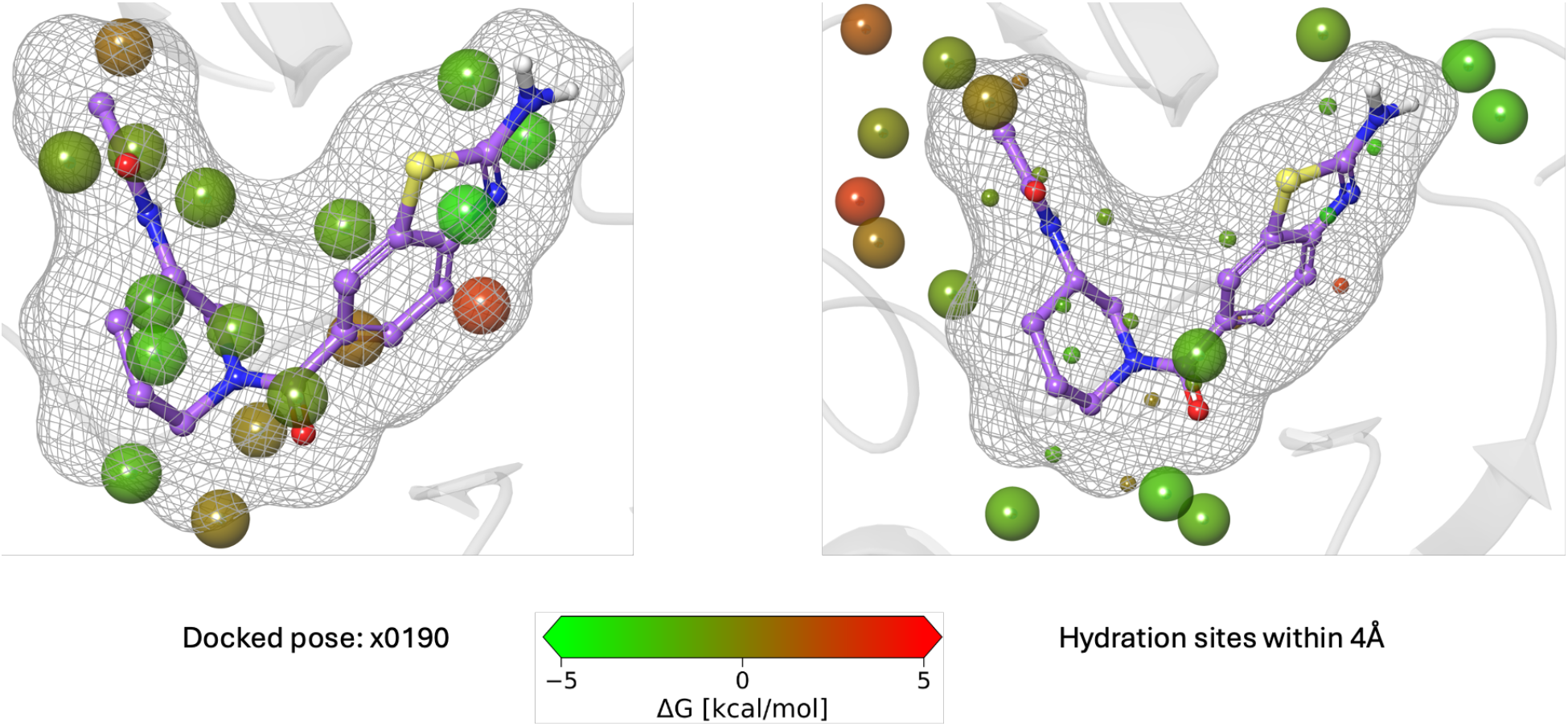
The WaterMap analysis of the docked complex of x0190 bound to ChikV Mac1.

In conclusion, x0190 and its analogs are promising leads for further exploration, most importantly by Mac1 inhibition assays in a wet lab^11^.

## Acknowledgments

This work was funded by the National Institute of Health (NIH) grant 1U19AI171110 to A.S. and I.E.

We thank Dr. Benjamin M. Webb for his help with our computing resources; Dr. Webb is a Nebius Fellow.

