## Supplementary material for "*In silico* analysis of the Chikungunya virus and SARS-CoV-2 Macrodomain": Supplementary_In silico analysis of the Chikungunya virus and SARS-CoV-2 Macrodomain.pdf

### Supplementary Figures and Tables

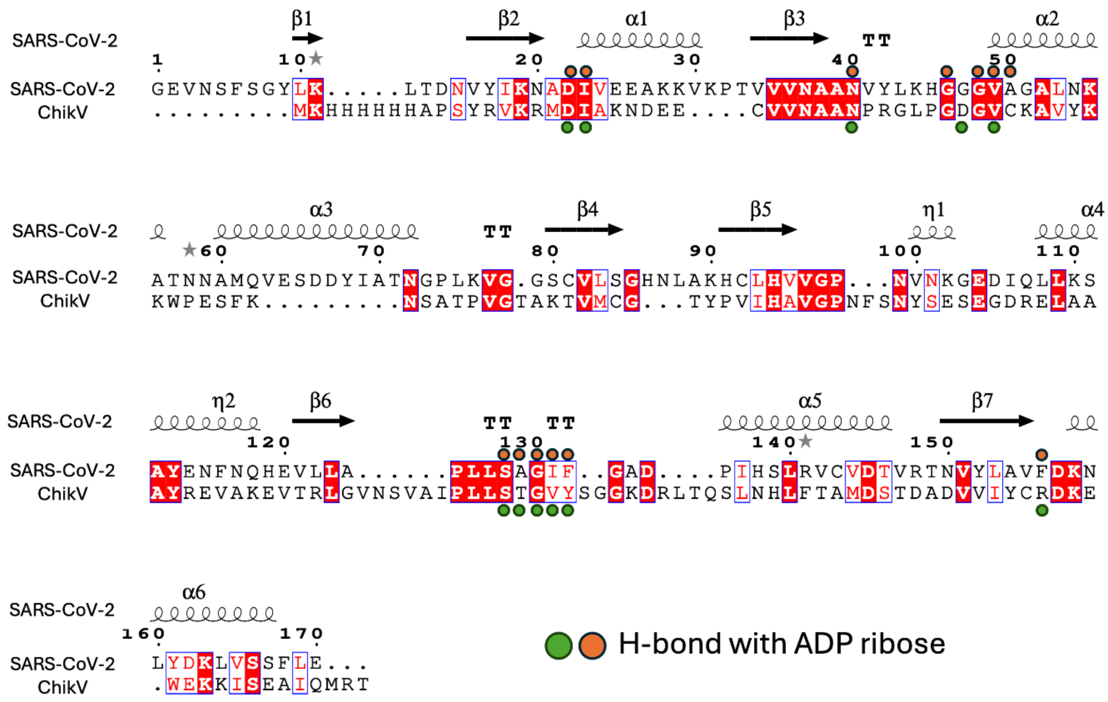


**Supplementary Figure 1:** Sequence alignment of SARS-CoV-2 and ChiKV Mac1. The residues that form H-bond with ADP ribose are marked with filled circles.

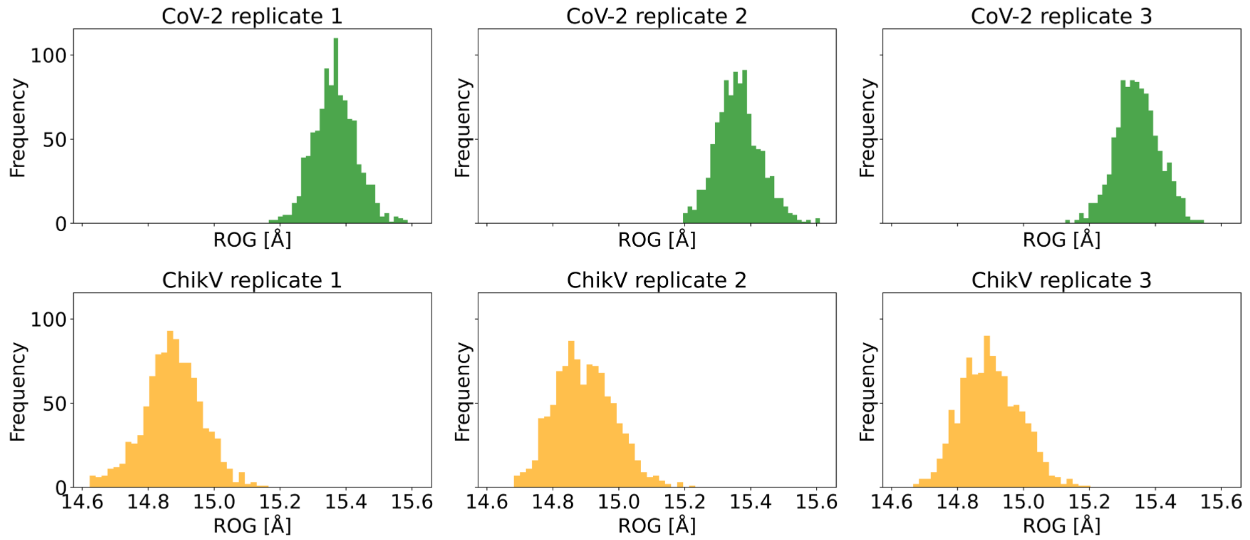


**Supplementary Figure 2:** Radius of gyration (ROG) calculated across the trajectory for apo ChikV and SARS-CoV-2 Mac1.

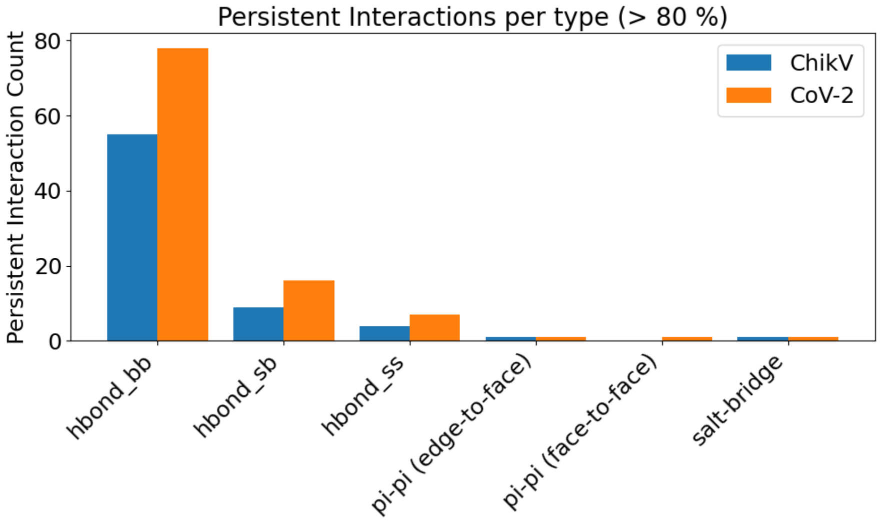


**Supplementary Figure 3:** Interaction persistence of different intra-protein non-covalent interactions in the apo Mac1 proteins across the simulation time. All three simulation trajectories were considered and interactions that persist for more than 80% of overall simulation time (3000 ns) were counted.

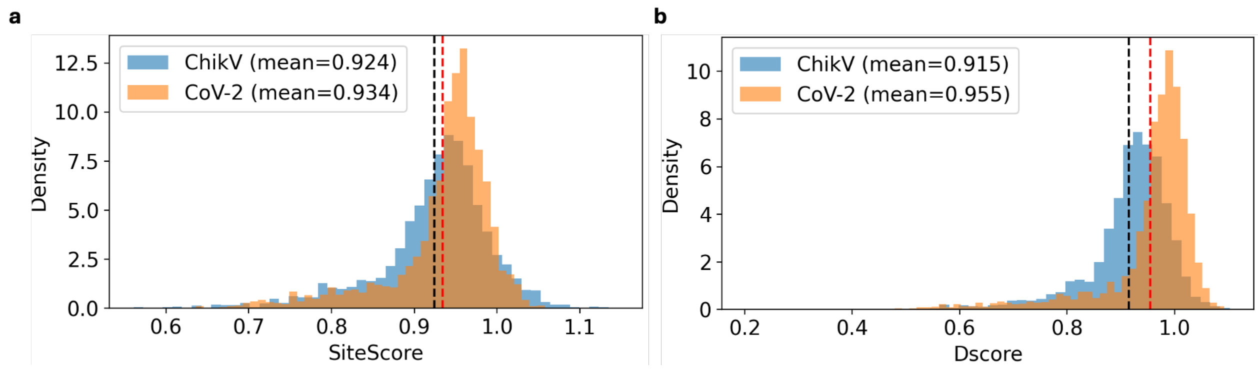


**Supplementary Figure 4:** ADP-ribose binding site analysis across trajectory for apo ChikV and SARS-CoV-2 Mac1. The SiteMap was run for each frame of all replicates and SiteScore and Dscore were calculated.

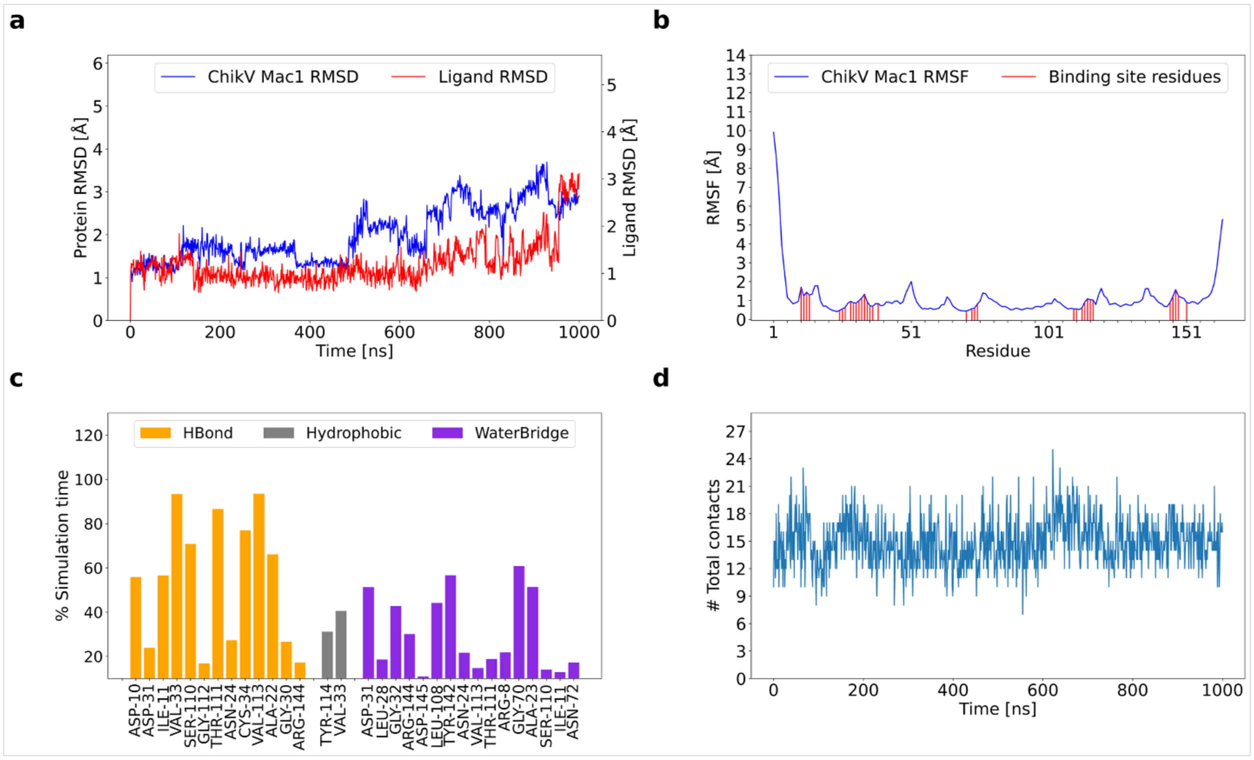


**Supplementary Figure 5:** MD simulation of ADP-ribose bound ChikV Mac1 (Replicate 2) (a) RMSD plot of Mac1 and ADP-ribose (ligand RMSD) (b) RMSF of ChikV Mac1. The ADP-ribose interacting residues are marked (c) Non-covalent interaction between ADP-ribose and ChikV Mac1. The y-axis represents the persistence of each interaction type. transient interactions (<10% persistence) have not been shown. (d) Total number of contacts of ADP-ribose across the simulation time.

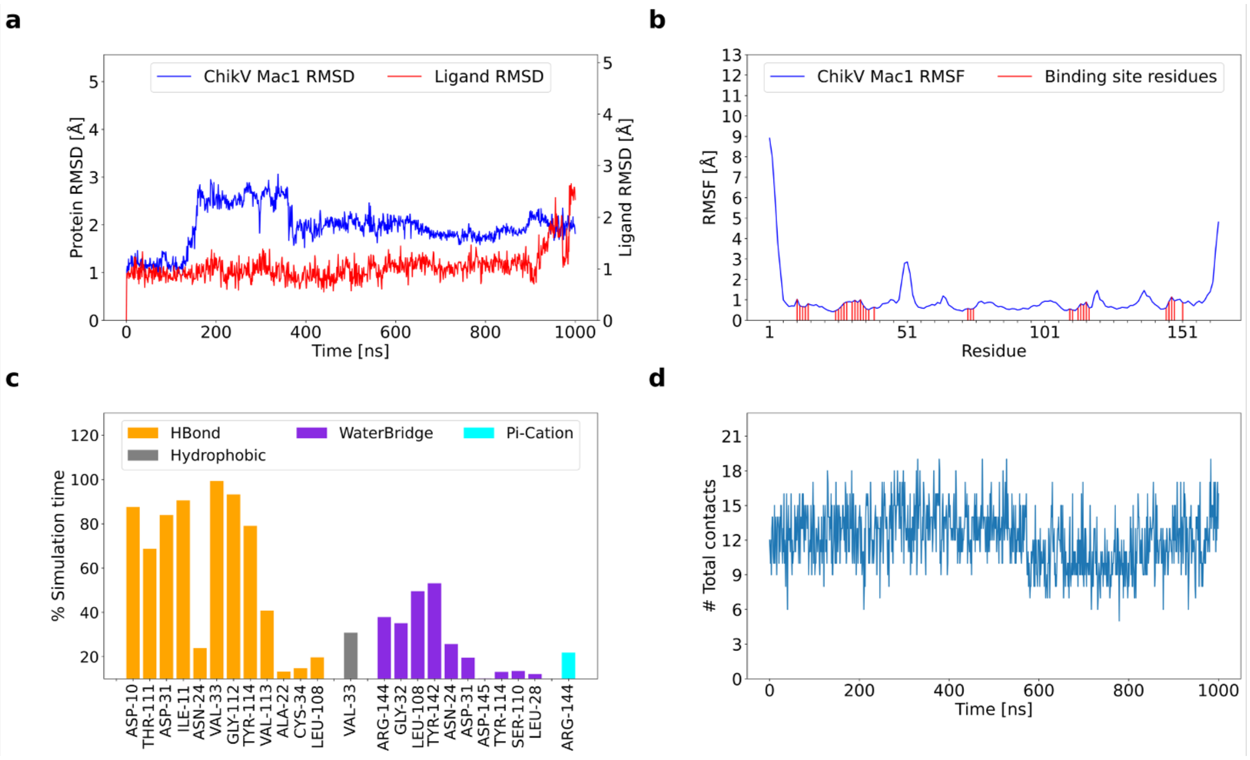


**Supplementary Figure 6:** MD simulation of ADP-ribose bound ChikV Mac1 (Replicate 3) (a) RMSD plot of Mac1 and ADP-ribose (ligand RMSD) (b) RMSF of ChikV Mac1. The ADP-ribose interacting residues are marked (c) Non-covalent interaction between ADP-ribose and ChikV Mac1. The y-axis represents the persistence of each interaction type. Transient interactions (<10% persistence) have not been shown. (d) Total number of contacts of ADP-ribose across the simulation time.

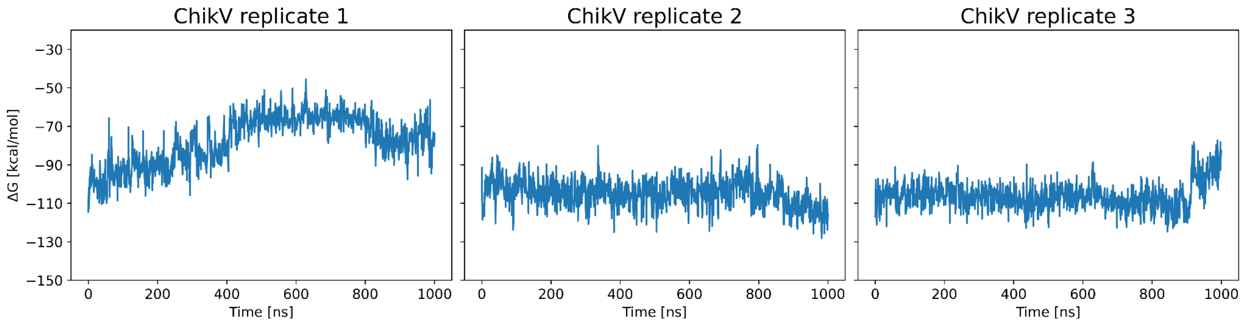


**Supplementary Figure 7:** The binding energy (ΔG) of ADP-ribose to ChikV Mac1 across the simulation time in all three replicates.

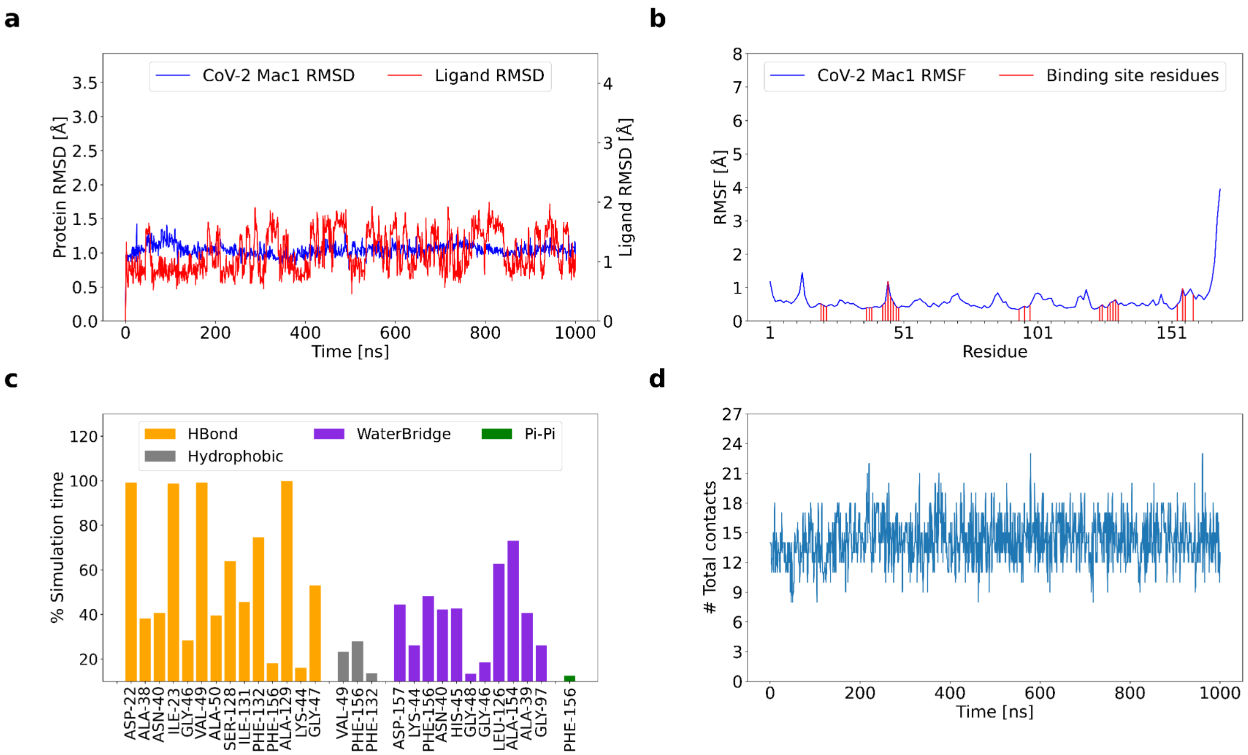


**Supplementary Figure 8:** MD simulation of ADP-ribose bound SARS-CoV-2 Mac1 (Replicate 1) (a) RMSD plot of Mac1 and ADP-ribose (ligand RMSD) (b) RMSF of SARS-CoV-2 Mac1. The ADP-ribose interacting residues are marked (c) Non-covalent interaction between ADP-ribose and SARS-CoV-2 Mac1. The y-axis represents the persistence of each interaction type. transient interactions (<10% persistence) have not been shown. (d) Total number of contacts of ADP-ribose across the simulation time.

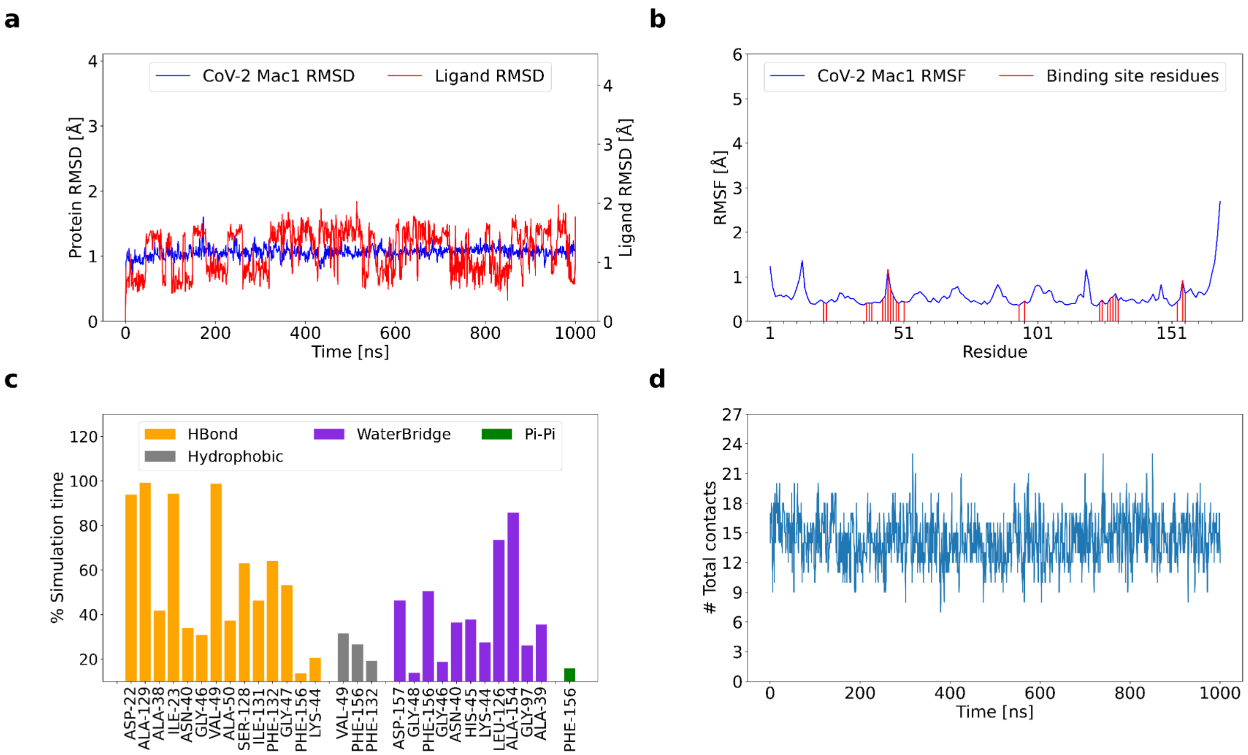


**Supplementary Figure 9:** Molecular dynamics simulations of ADP-ribose bound SARS-CoV-2 Mac1 (Replicate 2) (a) RMSD plot of Mac1 and ADP-ribose (ligand RMSD) (b) RMSF of SARS-CoV-2 Mac1. The ADP-ribose interacting residues are marked (c) Non-covalent interaction between ADP-ribose and SARS-CoV-2 Mac1. The y-axis represents the persistence of each interaction type. transient interactions (<10% persistence) have not been shown. (d) Total number of contacts of ADP-ribose across the simulation time.

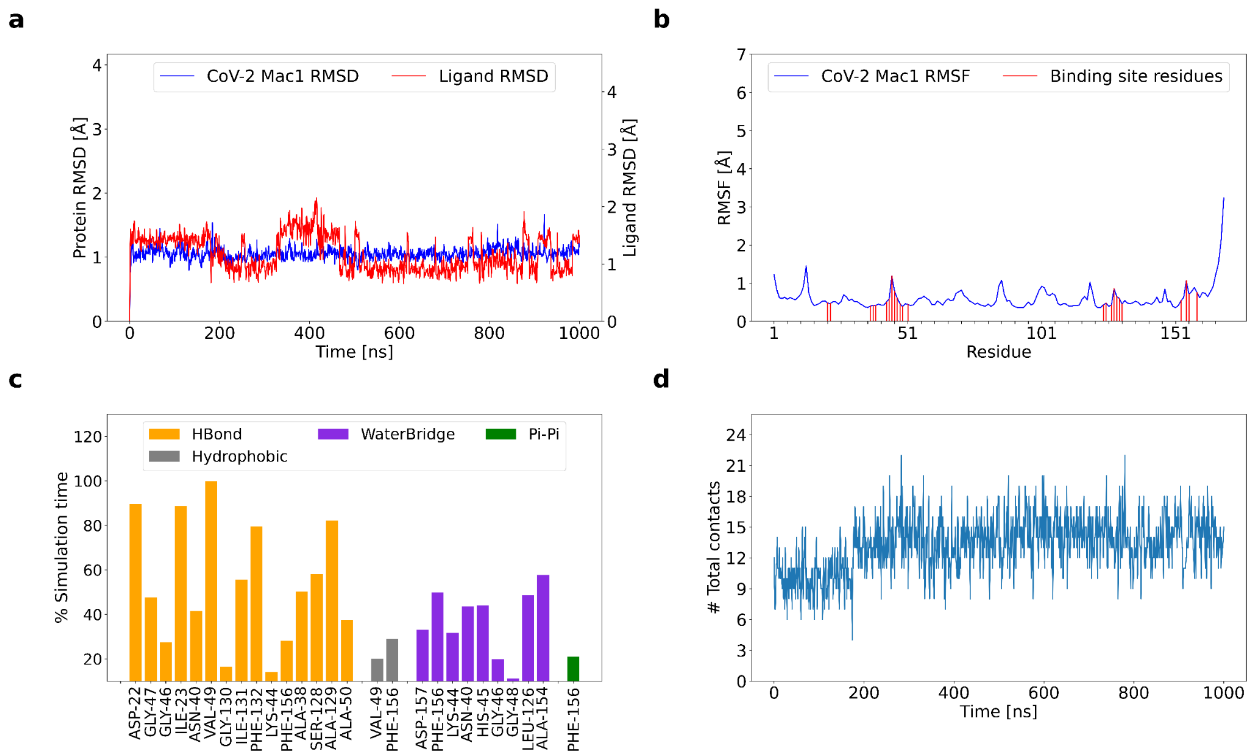


**Supplementary Figure 10:** Molecular dynamics simulations of ADP-ribose bound SARS-CoV-2 Mac1 (Replicate 3) (a) RMSD plot of Mac1 and ADP-ribose (ligand RMSD) (b) RMSF of SARS-CoV-2 Mac1. The ADP-ribose interacting residues are marked (c) Non-covalent interaction between ADP-ribose and SARS-CoV-2 Mac1. The y-axis represents the persistence of each interaction type. transient interactions (<10% persistence) have not been shown. (d) Total number of contacts of ADP-ribose across the simulation time.

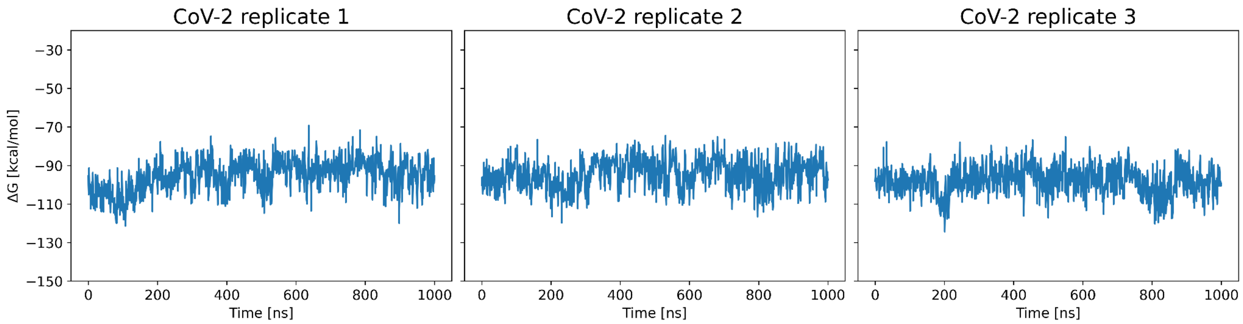


**Supplementary Figure 11:** The binding free energy (ΔG) of ADP-ribose to SARS-CoV-2 Mac1 across the simulation time in all three replicates.

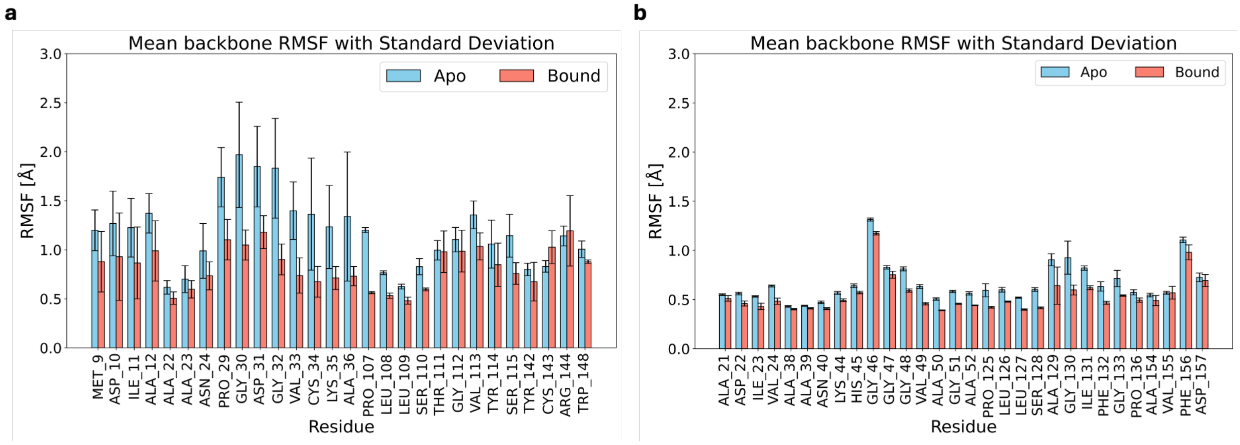


**Supplementary Figure 12:** The mean backbone RMSF of apo and holo Mac1. (a) ChikV. (b) SARS-CoV-2. The mean is calculated from all three trajectories.

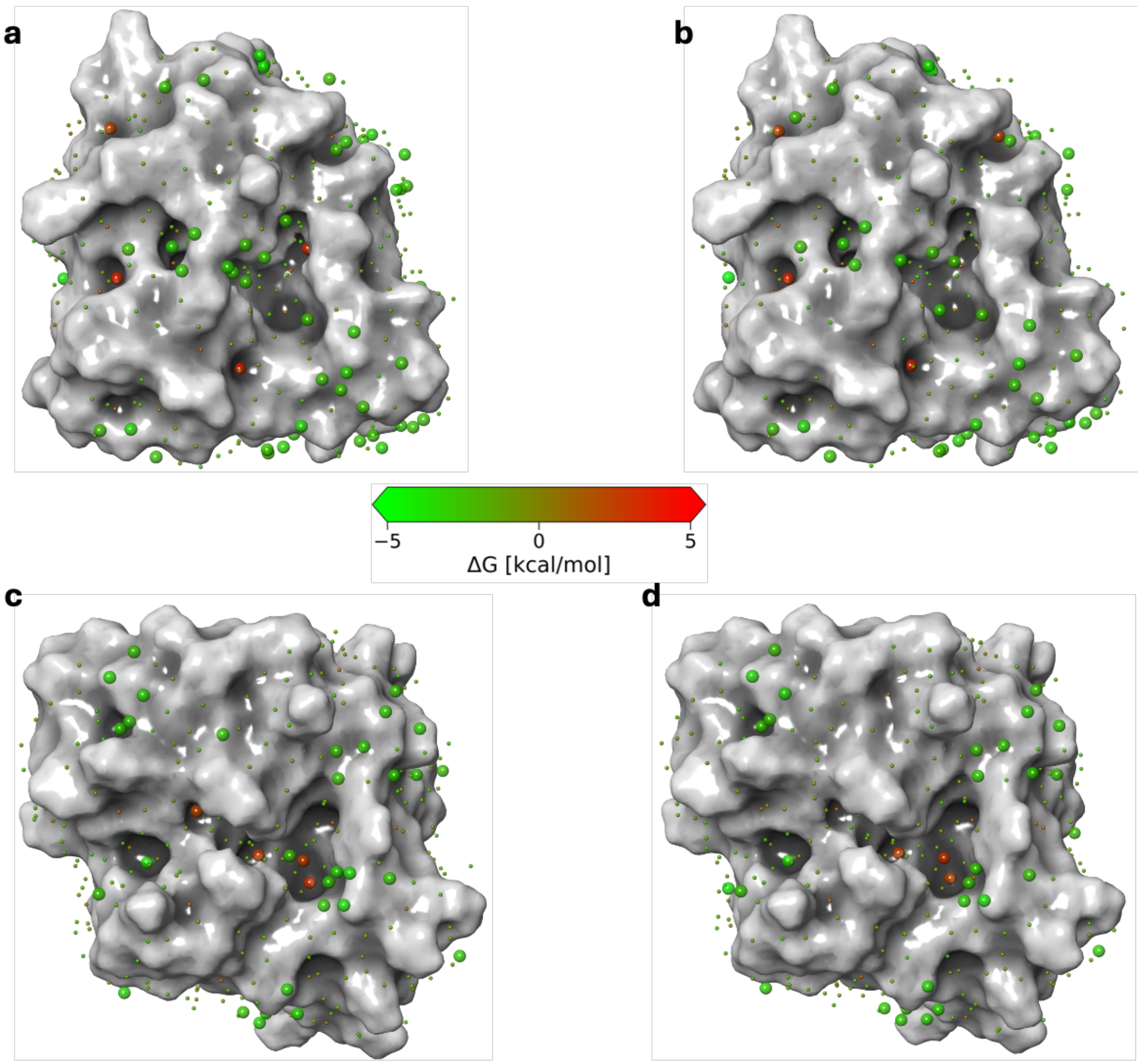


**Supplementary Figure 13:** WaterMap analysis of apo Mac1. (a, b) ChikV Mac1 hydration sites for replicates 2 and 3. (c, d) SARS-CoV-2 Mac1 hydration sites for replicates 2 and 3.

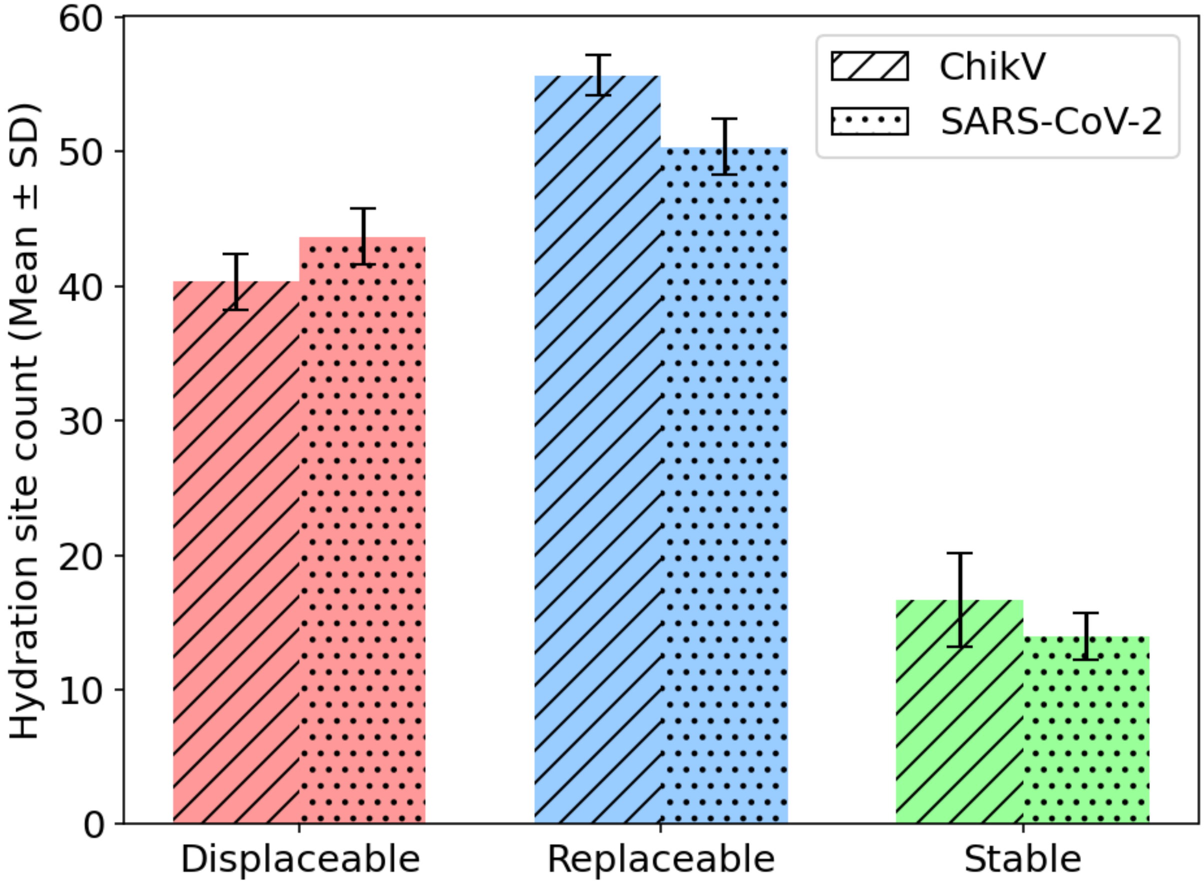


**Supplementary Figure 14:** Categorization of hydration sites within 4 Å of ADP-ribose binding site.

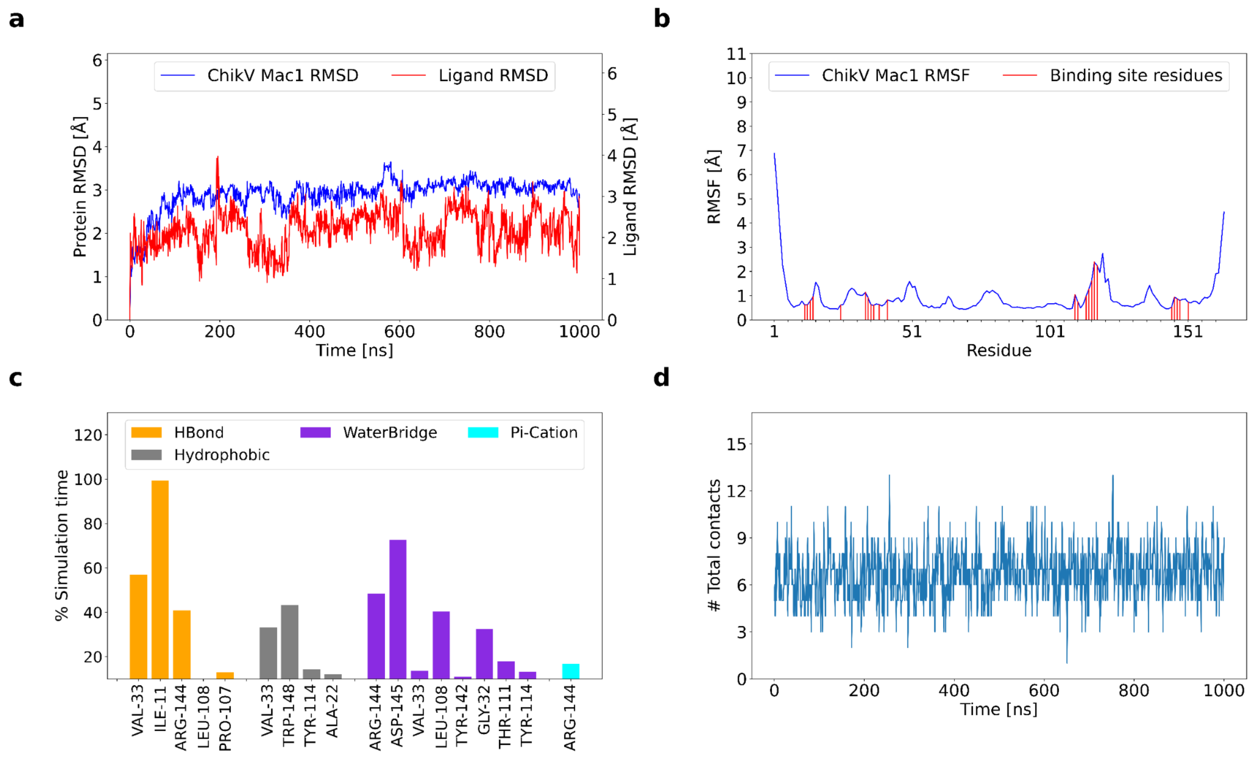


**Supplementary Figure 15:** Molecular dynamics simulations of x0190 compound bound to ChikV Mac1 (Replicate 1). (a) RMSD plot of Mac1 and x0190 (ligand RMSD). (b) RMSF of ChikV Mac1. The x0190 interacting residues are marked (c) Non-covalent interaction between x0190 and ChikV Mac1. The y-axis represents the persistence of each interaction type. Transient interactions (<10% persistence) have not been shown. (d) Total number of contacts of x0190 across the simulation time.

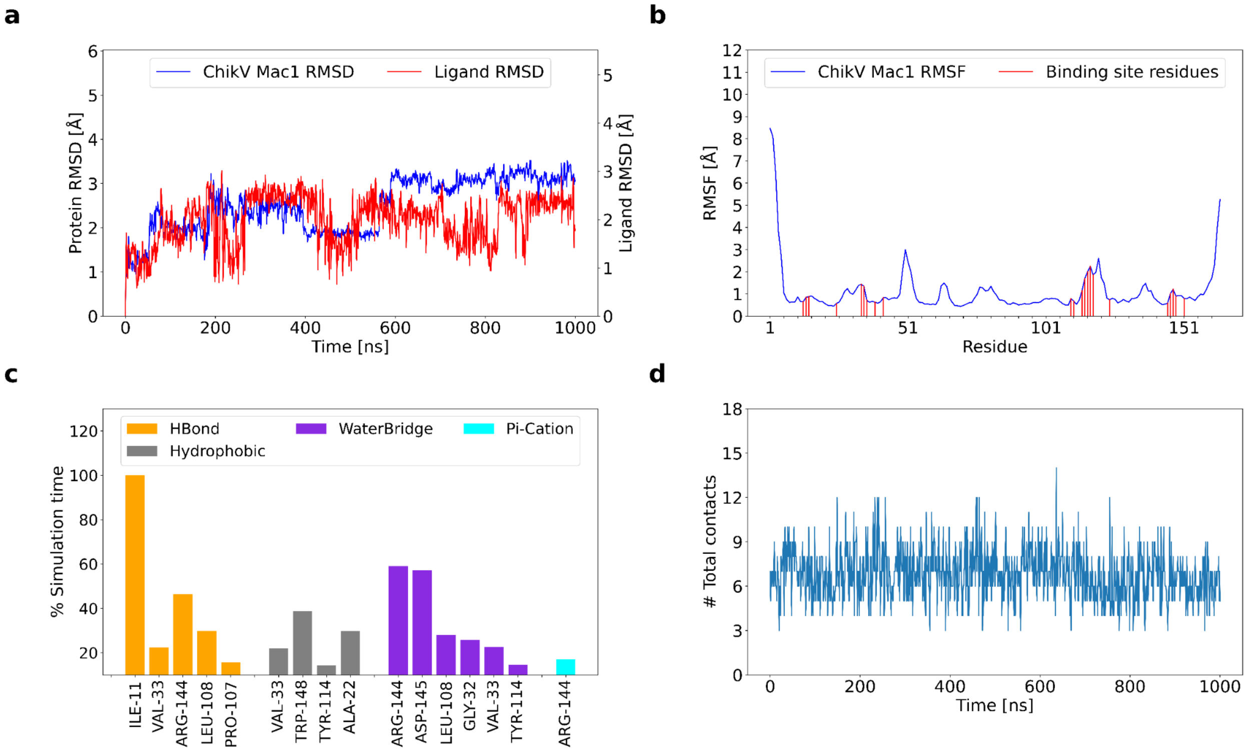


**Supplementary Figure 16:** Molecular dynamics simulations of x0190 compound bound to ChikV Mac1 (Replicate 1). (a) RMSD plot of Mac1 and x0190 (ligand RMSD). (b) RMSF of ChikV Mac1. The x0190 interacting residues are marked. (c) Non-covalent interaction between x0190 and ChikV Mac1. The y-axis represents the persistence of each interaction type. Transient interactions (<10% persistence) have not been shown. (d) Total number of contacts of x0190 across the simulation time.

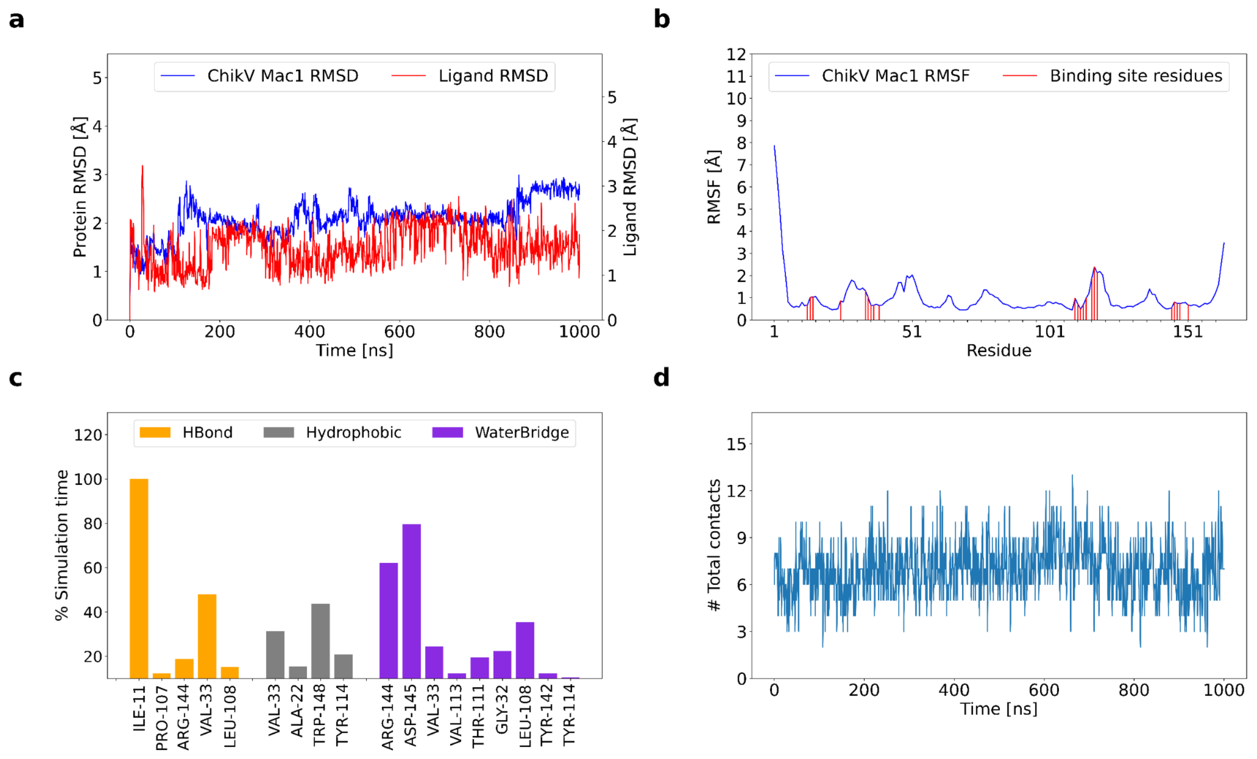


**Supplementary Figure 17:** Molecular dynamics simulations of x0190 compound bound to ChikV Mac1 (Replicate 1). (a) RMSD plot of Mac1 and x0190 (ligand RMSD). (b) RMSF of ChikV Mac1. The x0190 interacting residues are marked. (c) Non-covalent interaction between x0190 and ChikV Mac1. The y-axis represents the persistence of each interaction type. Transient interactions (<10% persistence) have not been shown. (d) Total number of contacts of x0190 across the simulation time.

**Supplementary Table 1:** Interactions of water molecules in hydration sites with Mac1 residues.

| Mac1 protein | Residues involved in H-bond interaction with stable hydration sites | Residues involved in H-bond interaction with replaceable hydration sites (ΔG>0 and ΔH<0) | Residues involved in H-bond interaction with displaceable hydration sites (ΔG>0 and ΔH>0) |
|---|---|---|---|
| ChikV (Replicate 1) | Asp10, Asp31, Val33, Thr111 | Met9, Asp10, Ile11, Ala23, Gly30, Asp31, Gly32, Val33, Lys35, Ala36, Leu108, Ser110, Thr111, Gly112, Val113, Tyr114, Ser115, Tyr142, Cys143, Arg144, Trp148 | Asp10, Ala12, Ala22, Ala23, Asn24, Asp31, Gly32, Cys34, Lys35, Leu108, Thr111, Gly112, Val113, Tyr114, Tyr142, Arg144 and Trp148 |
| ChikV (Replicate 2) | Asp10, Asp31, Gly32, Val113 | Asp10, Ile11, Ala23, Gly30, Asp31, Gly32, Val33, Lys35, Ala36, Leu108, Ser110, Thr111, Gly112, Val113, Ser115, Tyr142, Arg144 Trp148 | Met9, Asp10, Ile11, Ala12, Ala22, Ala23, Asn24, Gly30, Asp31, Gly32, Cys34, Lys35, Leu108, Thr111, Gly112, Val113, Tyr114, Ser115, Tyr142, Cys143, Arg144 |
| ChikV (Replicate 3) | Asp31, Thr111, Val113 | Met9, Asp10, Ile11, Ala22, Ala23, Gly30, Asp31, Gly32, Val33, Cys34, Lys35, Ala36, Leu108, Thr111, | Asp10, Ala12, Ala22, Ala23, Asn24, Gly30, Asp31, Gly32, Lys35, Leu108, Ser110, Thr111, Gly112, |

| | | | |
|---|---|---|---|
| | | Gly112, Val113, Tyr114, Ser115, Tyr142, Arg144, Trp148 | Val113, Tyr114, Ser115, Tyr142, Arg144 |
| SARS-CoV-2 (Replicate 1) | Asp226 and His249 | Asp226, Ala242, Ala243, Asn244, Lys248, His249, Gly250, Gly251, Gly252, Val253, Ala254, Gly255, Asn303, Ala333, Gly334, Ile335, Gly337 and Phe360 | Ala225, Ile227, Ala242, Asn244, Lys248, His249, Gly250, Gly251, Gly255, Ala256, Asn303, Leu330, Ser332, Ala333, Gly334, Ile335, Gly337, Ala358, and Phe360 |
| SARS-CoV-2 (Replicate 2) | Asp226, Lys248 and His249 | Asp226, Ala242, Ala243, Asn244, Lys248, His249, Gly250, Gly251, Gly252, Val253, Ala254, Gly255, Asn303, Ala333, Gly334, Ile335, Gly337 and Phe360 | Ala225, Ile227, Ala242, Asn244, Lys248, His249, Gly250, Gly252 Gly255, Ala256, Asn303, Leu330, Ser332, Ala333, Gly334, Ile335, Gly337, Ala358, and Phe360 |
| SARS-CoV-2 (Replicate 3) | Asp226 and Lys248 | Asp226, Ala242, Ala243, Asn244, Lys248, His249, Gly250, Gly251, Gly252, Val253, Ala254, Gly255, | Ala225, Ile227, Ala242, Asn244, Lys248, His249, Gly250, Gly252 Gly255, Ala256, Asn303, Leu330, |

| | | | |
|---|---|---|---|
| | | Asn303, Gly334, Ile335, Gly337, Ala358 and Phe360 | Ser332, Ala333, Gly334, Ile335, Gly337, Ala358, Phe360 and Leu364 |
